# Duplex perception reveals brainstem auditory representations are modulated by listeners’ ongoing percept for speech

**DOI:** 10.1101/2023.05.09.540018

**Authors:** Rose Rizzi, Gavin M. Bidelman

**Affiliations:** Department of Speech, Language, and Hearing Sciences, Indiana University, Bloomington, IN, USA; Program in Neuroscience, Indiana University, Bloomington, IN, USA; Cognitive Science Program, Indiana University, Bloomington, IN, USA; School of Communication Sciences and Disorders, University of Memphis, Memphis, TN, USA

**Author notes:** Author contributions: G.M.B. designed the experiment, R.R. collected the data, R.R. and G.M.B. analyzed the data and wrote the paper. Address for editorial correspondence: Gavin M. Bidelman, Ph.D., Speech, Language and Hearing Sciences 2631 East Discovery Parkway Bloomington, IN 47408.

**Keywords:** Brainstem response, binaural processing, categorical perception, electroencephalography (EEG), frequency-following response (FFR)

## Abstract

So-called duplex speech stimuli with perceptually ambiguous spectral cues to one ear and isolated low– vs. high-frequency third formant “chirp” to the opposite ear yield a coherent percept supporting their phonetic categorization. Critically, such dichotic sounds are only perceived categorically upon binaural integration. Here, we used frequency-following responses (FFRs), scalp-recorded potentials reflecting phase-locked subcortical activity, to investigate brainstem responses to fused speech percepts and to determine whether FFRs reflect binaurally integrated category-level representations. We recorded FFRs to diotic and dichotic stop-consonants (/da/, /ga/) that either did or did not require binaural fusion to properly label along with perceptually ambiguous sounds without clear phonetic identity. Behaviorally, listeners showed clear categorization of dichotic speech tokens confirming they were heard with a fused, phonetic percept. Neurally, we found FFRs were stronger for categorically perceived speech relative to category-ambiguous tokens but also differentiated phonetic categories for both diotically and dichotically presented speech sounds. Correlations between neural and behavioral data further showed FFR latency predicted the degree to which listeners labeled tokens as “da” vs. “ga”. The presence of binaurally integrated, category-level information in FFRs suggests human brainstem processing reflects a surprisingly abstract level of the speech code typically circumscribed to much later cortical processing.

---

Listeners effortlessly use continuous acoustic information in the soundscape to form perceptual categories that enable speech perception (Liberman et al. 1957; Liberman et al. 1967; Pisoni 1973). Such categorical percepts remain relatively invariant to acoustic changes within category, which may ultimately help listeners cope with challenges to the speech signal including speaker variability (Sumner 2011) or background noise (Bidelman et al. 2020; Carter and Bidelman 2021). The highly categorical nature of speech has led some to suggest it is heard via a specialized “phonetic mode” of listening (Mann and Liberman 1983; Whalen and Liberman 1987; Liberman and Mattingly 1989). Still, it is now clear categories are not unique to speech, *per se,* but extend to a variety of cognitive processes such as face (Beale and Keil 1995), color (Franklin et al. 2008) and music perception (Burns and Ward 1978; Zatorre 1983; Mankel et al. 2022). Though categorization is central to our understanding of speech processing, the neural mechanisms for this perceptual phenomenon remain controversial.

Because phonetic labeling is a broad cognitive process, studies on the neural underpinnings of auditory category representation have focused nearly exclusively on *cortical* mechanisms (Maiste et al. 1995; Sharma and Dorman 1999; Chang et al. 2010; Bidelman et al. 2013; Carter and Bidelman 2021). Despite this focus on category perception as a cortical process, recent evidence suggests category-level information might arise prior to neocortex. In this regard, the frequency-following response (FFR) has been a useful tool for examining subcortical auditory processing and the neural encoding of pitch, timbre, and timing elements of speech (Galbraith et al. 1995; Krishnan 2002; Skoe and Kraus 2010; Bidelman and Powers 2018). More recent FFR studies have shed new light on the emergence of subcortical category representations (Carter and Bidelman 2023). FFRs are scalp-recorded potentials reflecting a mixture of phase locked activity from several nuclei along the auditory pathway (Smith et al. 1975; Sohmer et al. 1977; Skoe and Kraus 2010; Bidelman 2015; Bidelman 2018). Though cortex can contribute to FFRs under limited circumstances (Coffey, Herholz, et al. 2016), speech-FFRs are dominantly generated by midbrain sources (i.e., inferior colliculus, IC) when recorded via EEG using high (> 150 Hz) fundamental frequency stimuli (Kiren et al. 1994; Bidelman 2018; López-Caballero et al. 2020; Bidelman and Momtaz 2021; Gorina-Careta et al. 2021; Price and Bidelman 2021). While the IC is likely too early along the processing hierarchy to show bottom-up, categorical organization *de novo*, top-down influences from cortex via the descending corticofugal system (Gao and Suga 1998) could modulate brainstem speech representations to produce categorical encoding effects as was recently observed in human FFRs (Price and Bidelman 2021; Lai et al. 2022; Carter and Bidelman 2023; Lai et al. 2023). Indeed, categorical coding in FFRs is not observed under passive listening (Carter and Bidelman 2023) suggesting goal-directed attention is necessary for the corticofugal system to exert real-time influences on midbrain speech processing (Lai *et al*. 2022).

Related controversy surrounds the FFR and whether it reflects a true *perceptual* correlate or simply a *neuro-acoustic* representation of complex sounds (Gockel et al. 2011; Bidelman *et al*. 2013; Coffey, Colagrosso, et al. 2016; Yellamsetty and Bidelman 2019; Carter and Bidelman 2023). Speech-FFRs follow the time-frequency cues of speech with remarkably fidelity to the point they are intelligible to listeners when sonified (i.e., replayed) as an audio stimulus (Galbraith *et al*. 1995; Weiss and Bidelman 2015; Bidelman 2018). Still, because the FFR is neurophonic, changes in acoustic stimulus properties will produce corresponding changes in the neural response. This conflation of variables with behavior makes it difficult to isolate whether differences in neural activity truly index perception (endogenous coding) or are instead due to trivial acoustic mirroring (exogenous coding) (Carlyon 2004). Still, converging evidence suggests perceptual information may indeed drive changes in FFR when stimuli are well controlled and pit acoustic information orthogonal to the resulting percept—as afforded by categorization tasks (Price and Bidelman 2021; Lai *et al*. 2022; Carter and Bidelman 2023; Lai *et al*. 2023). Additionally, speech percepts that require binaural integration between the ears offer another viable test of perceptual correlates in FFR because information must be fused centrally to generate a category label. Indeed, there is already some evidence that the FFR reflects binaural auditory percepts (Galbraith et al. 1998; Krishnan and McDaniel 1998; Bidelman and Krishnan 2009). Here, we extend these ideas by using stimuli which support “duplex” speech perception (Preisig and Sjerps 2019) to further probe phonetic-level coding in FFR.

Duplex perception refers to the binaurally fused categorical percept of dichotic stop-consonant stimuli in which an ambiguous portion of the spectrum (i.e., low-frequency “base”) is presented to one ear while a disambiguating portion of the spectrum containing the third formant (F3) (i.e., high-frequency “chirp”) is presented to the other (Rand 1974; Liberman et al. 1981; Mann and Liberman 1983; Preisig and Sjerps 2019). Critically, the cues at each individual ear are phonetically ambiguous. However, when heard together, duplex stimuli are perceived as a fused speech percept with a clear category label; varying the frequency of the F3 chirp produces percepts from /ga/ (low F3) to /da/ (high F3). Because these stimuli share an identical spectral base presented to one ear with acoustic information varying only in F3 at the other, their spectral content is uniquely controlled. Yet, they support category labeling through binaural integration. While neuroimaging studies have investigated duplex speech perception using cortical EEG (Gokcen and Fox 2001; Pérez et al. 2008) and fMRI (Preisig and Sjerps 2019; Preisig et al. 2020; Preisig et al. 2021; Preisig et al. 2022), we are aware of no studies that have used these binaurally fused sounds to investigate speech encoding at the brainstem level. This approach would allow a novel test of whether FFRs carry a higher-level, perceptual correlate of speech beyond its acoustic features.

To this end, the current study aimed to evaluate whether speech-FFRs are modulated by listeners’ perception, reflect binaural integration, and carry category level information of the speech signal. We measured FFRs in response to dichotic duplex and diotic speech stimuli in younger adults during a novel categorization paradigm task that allows for simultaneous recording of brainstem responses during real time behavior (e.g., Bidelman 2015; Carter and Bidelman 2023). Critically, the high-frequency bandwidth of our stimuli (> 250 Hz) was designed to far exceed the low-frequency (< 100 Hz) phase-locking capacity of cortical neurons (Joris et al. 2004) and thus ensure our FFRs were of a subcortical origin (Brugge et al. 2009; Bidelman 2018; Gorina-Careta *et al*. 2021). Based on previous literature suggesting speech-FFRs are influenced by categories, attention, and listening experience, we hypothesized that responses to duplex stimuli would more closely mirror phonetic rather than acoustic dimensions of the stimulus. Our findings reveal that both the strength and timing of the FFR are modulated by listeners’ ongoing phonetic percepts, reflecting binaural integration and category level representations of the speech signal.

## Materials and Methods

### Participants

The sample included *N*=16 monolingual English-speaking young adults (age range=22-28 years,15 female) with an average of 18.25 ± 1.29 years of education. Participants all had normal hearing (pure tone thresholds ≤25 dB HL; 250-8000 Hz), had varied musical experience (mean=5.56 ± 6.4 years), and were mostly right-handed (mean=76% ± 27%; Edinburgh Handedness Inventory; Oldfield 1971). Each participant provided written consent in compliance with a protocol approved by the Institutional Review Board of The University of Memphis. Participants were paid $10 an hour for their time.

### Stimuli and task

Stimuli consisted of synthetic /da/ and /ga/ consonant vowel (CV) speech tokens that were presented diotically or dichotically (duplex stimuli) to listeners (**Figure 1**) (stimuli acquired from: https://asa.scitation.org/doi/suppl/10.1121/1.5092829) (Preisig and Sjerps 2019; Preisig *et al*. 2020; Preisig *et al*. 2021). The fundamental frequency (F0) of each speech token was 247 Hz, which is well above phase-locking limits of cortical neurons and thus ensured FFRs were of a subcortical origin (Joris *et al*. 2004; Brugge *et al*. 2009; Bidelman 2018; Gorina-Careta *et al*. 2021). Duplex stimuli were composed of an ambiguous base delivered to the right ear and a chirp (isolated 3^rd^ formant frequency; F3) delivered to the left ear (dichotic presentation). The base contained spectral information for F1, F2, and F4 formant frequencies of an /a/ vowel. Isolated chirps contained either a high (∼2.9 kHz) or low (∼2.7 kHz) F3 contour, promoting a /da/ or /ga/ percept, respectively. Critically, these dichotic stimuli require listeners to combine cues from both ears through binaural integration to properly arrive at a categorical label (i.e., “da” vs. “ga”); they cannot be classified via a single ear alone. In addition to duplex tokens, the base and chirp were presented diotically as control conditions. Listeners easily classify these latter tokens since the acoustic signal itself contains all category-relevant cues. The ambiguous base by itself served as an additional control. Each token was 160 ms in duration and tokens were gated with 5 ms ramps. In total, there were five stimulus conditions: ambiguous base + high F3 (promoting the percept of “da”), ambiguous base + low F3 (promoting the percept of “ga”), diotic /da/, diotic /ga/, and the ambiguous base alone.

**Figure 1:**
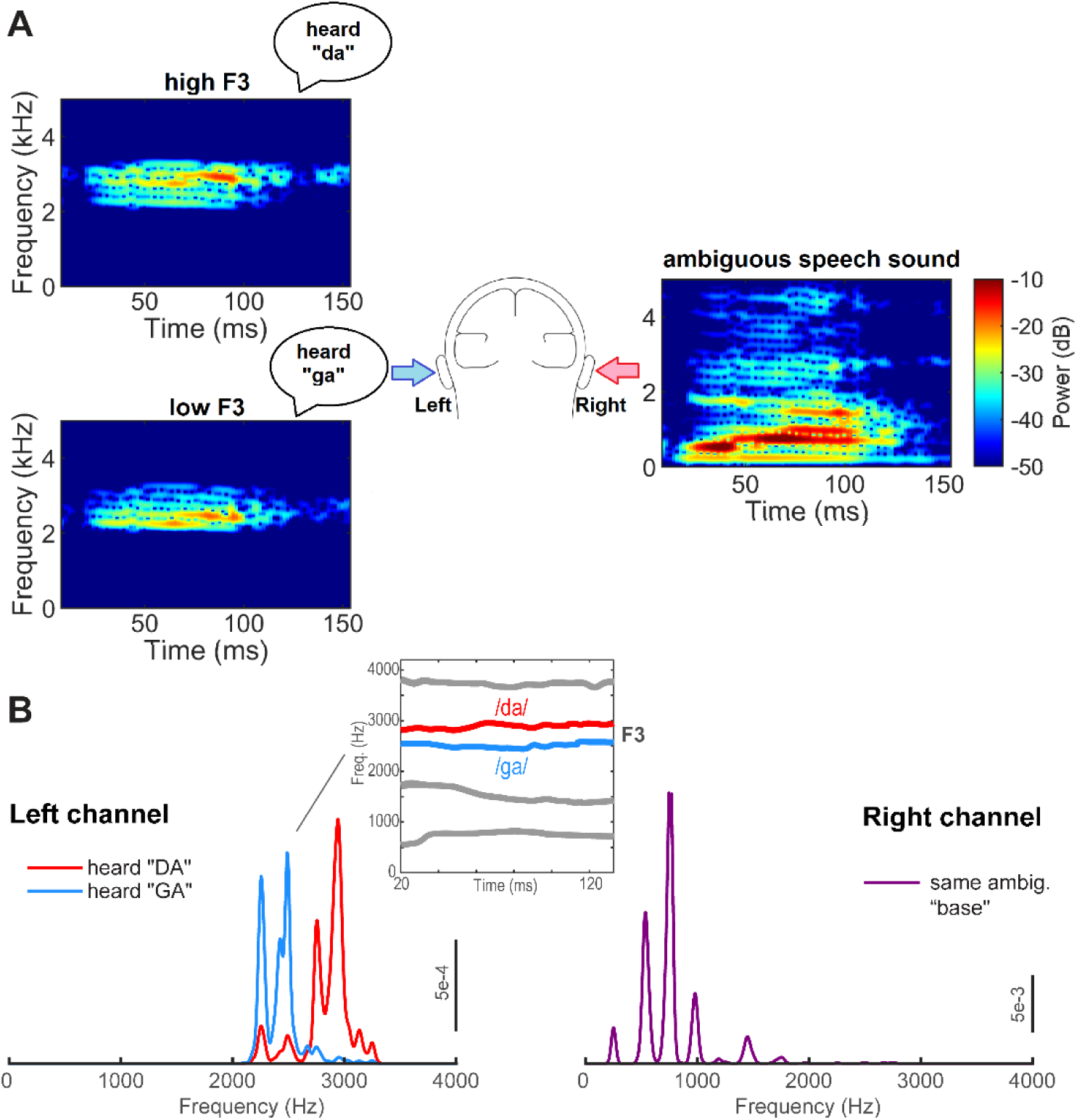
Duplex speech stimuli. (**A**) Spectrograms for F3 chirp and ambiguous base stimuli presented to each ear. (**B**) Spectra for duplex stimuli with F3 chirps in the left channel and ambiguous base in the right channel. The inset shows time-varying formant tracts for /da/ and /ga/ stimuli.

To efficiently record FFRs during an online behavioral task while obtaining the high (i.e., several thousand) trial counts needed for response visualization, we used a clustered interstimulus interval (ISI) presentation paradigm (Bidelman 2015; Carter and Bidelman 2023). Single tokens were presented in blocks of 30 repetitions with a rapid ISI (10 ms). After the clustered block of tokens ended, the ISI was slowed to 300 ms and a single token was presented to cue the behavioral response. Participants then indicated their percept (/da/ or /ga/) as quickly and accurately as possible via the keyboard. Following the behavioral response and a period of silence (250 ms) the next trial cluster commenced. This paradigm allowed 1980 presentations per token for input to FFR analysis and 66 tokens for the behavioral response.

Stimulus presentation was controlled via MATLAB (The MathWorks, Natick, MA, USA) routed to a TDT RP2 (Tucker-Davis Technologies, Alachua, FL, USA) signal processor. Stimuli were presented binaurally at 80 dB SPL via shielded insert headphones (ER-2; Etymotic Research) to prevent pickup of electromagnetic artifacts from contaminating neural recordings (Campbell et al. 2012; Price and Bidelman 2021).

### FFR recording

During the categorization task, continuous EEG activity was recorded differentially between scalp Ag/AgCl disk electrodes placed on the high forehead at the hairline (∼Fpz) referenced to linked mastoids (M1/M2); a mid-forehead electrode served as ground. This montage is optimal for pickup of the vertically oriented FFR dipole in the midbrain (Bidelman 2015). Electrode impedances remained ≤ 5 kΩ throughout the duration of recording. EEGs were digitized at 10 kHz to capture the fast activity of FFR. Raw EEG waveforms were epoched (0-165 ms), band-pass filtered (200-2500 Hz) to eradicate ocular artifacts and cortical activity thereby isolating brainstem responses (Musacchia et al. 2008; Bidelman *et al*. 2013), and averaged for each token per listener. Data preprocessing was performed in the MATLAB package Brainstorm (Tadel et al. 2011).

### Behavior data analysis

We calculated percent identification (percent of presentations of a token identified as /ga/) and reaction times (RTs) per stimulus condition. Improbable RTs (i.e., ≤ 250 ms or >2500 ms) were treated as fast guesses and lapses of attention, respectively, and were removed from the analysis as outliers (Bidelman and Walker 2017).

### FFR data analysis

We used Brainstorm to generate Fast Fourier Transforms (FFTs) for each token to quantify the spectral information in each response. We then measured the amplitude of the spectral peaks in a window of ±50 Hz around the nominal F0 (247 Hz). The F0 —related to voice pitch— represents the dominate energy in the FFR and is modulated by attention and listeners’ trial-by-trial categorical hearing (Price and Bidelman 2021; Lai *et al*. 2022; Carter and Bidelman 2023). We should note that the F0 used in the current study is considerably higher than those in nearly all previous work. Although our tokens all had identical voice pitch, we expected changes in F0 amplitude across tokens, indicating a modulation in the strength of the FFR dependent on listeners’ online percept (Carter and Bidelman 2023) and category cues integrated from the other ear. Onset latency was measured as the peak in the cross-correlation function between the FFR and evoking stimulus waveform in a 5.5-10 ms search window, the expected onset latency of the brainstem response (Galbraith and Brown 1990; Bidelman and Momtaz 2021).

### Statistical analysis

We used one-way mixed model ANOVAs (R; lme4 package, version 1.1-32) to analyze the FFR data (F0 amplitude and latency). The model included a fixed effect for token (5 levels: /da/, /ga/, duplex /da/, duplex /ga/, ambiguous base) and a random effect for subjects. Identical models were run for the behavioral measures (percent /ga/, RTs). We normalized the FFR amplitude measures between 0-1 (within each subject) to mirror the behavioral percent /ga/ identification data which is similarly bound between 0-100%. This allowed us to focus on the *relative* changes across stimulus conditions in both neural and behavior measures on similar scales. To assess relationships between perceptual and neural responses, we computed repeated measures correlations (rmCorr R package, version 0.5.4) (Bakdash and Marusich 2017) between FFR measures and identification scores. Unlike conventional correlations, repeated measures correlations account for non-independence among observations, adjusts for between subject variability, and measure within-subject correlations by evaluating the common intra-individual association between two measures.

### Simulated FFRs from a computational AN model

We next aimed to test whether our dichotically-evoked FFRs and perceptual correlates could be explained by a mere summation of responses to the other ear and thus reflect acoustic-rather than perceptually-based (binaurally integrated) coding. Binaural interaction is typically measured as the residual difference between the binaural and summed monaural responses [i.e., FFR_binaural_ – (FFR_LE_ + FFR_RE_) > 0] (Wernick and Starr 1968; Gerken et al. 1975; Krishnan and McDaniel 1998). However, as noted by Hink et al. (1980), unmeasured differences in signal-to-noise ratio of each ear’s monaural recordings can result in spurious estimates of binaural interaction. Additionally, the entirety of our chirp stimulus spectrum exceeds 2000 Hz (see Fig. 1), which is beyond the phase-locking capacity of FFRs (Bidelman and Powers 2018). Therefore, FFRs to the chirps alone could not be recorded.

To circumvent these confounds, we instead opted to use a computational model of the auditory nerve (AN) (Zilany et al. 2014) to simulate brainstem FFRs to dichotic stimuli. Details of this phenomenological AN model and FFR simulation are provided by Zilany et al. (2009) and Bidelman (2014), respectively. The model incorporates several important nonlinearities observed in the auditory periphery, including cochlear filtering, level-dependent gain (i.e., compression) and bandwidth control, long-term adaptation, as well as two-tone suppression. Model tuning curves were fit to the characteristic frequency (CF)-dependent variation in threshold and bandwidth for high-spontaneous rate (SR) fibers in normal-hearing cats (Miller et al. 1997). The stochastic nature of AN responses is accounted for by a modified non-homogenous Poisson process, which includes effects of both absolute and relative refractory periods and captures the major stochastic properties of single-unit AN responses (e.g., Young and Barta 1986).

We used the AN model to simulate scalp-recorded speech-FFRs (Bidelman 2014; Carter and Bidelman 2023) (**Fig. 6A**). This approach is based on the assumption that the far-field FFR recorded at the scalp is a convolution of an elementary unit waveform (i.e., impulse response; akin to the click-evoked ABR) with the instantaneous discharge rate from a given auditory nucleus (Goldstein and Kiang 1958; Dau 2003). The modeling pipeline was otherwise identical to Carter and Bidelman (2023) with the exception that we used the latest generation of the model that incorporates revised estimates of human cochlear tuning based on otoacoustic emission data (Shera et al. 2002).

We submitted 50 repetitions of each stimulus to the model to evoke AN spike-trains. Spikes were generated from each of 100 model fibers (CFs: 125-11000 Hz; high spontaneous rate units) to simulate the discharge pattern across the cochlear partition. Activity from the entire ensemble was then summed to form a population post-stimulus time histogram (PSTH). The PSTH was then convolved with a unitary response function, simulating the impulse response of nuclei from the auditory brainstem (for details, see Dau 2003). Finally, pink noise (1/*f* distribution) was added to simulate the quasi-stochastic nature of EEG noise (Granzow et al. 2001; Dau 2003; Bidelman 2014). Resulting model waveforms provided a close approximation of the time-frequency characteristics of true FFRs recorded in our human listeners.

To simulate dichotic FFRs, we generated model FFR outputs separately for the left and right audio channels of our duplex stimuli. We then summed the monaural FFRs to simulate binaurally fused responses as measured in the actual FFR experiment. As with the empirical FFR recordings, we then measured model F0 (247 Hz) amplitudes from response spectra. This allowed us to compare true FFR with model responses, which similarly reflect the output of cochlear processing (e.g., spectral decomposition, nonlinearities) but are not subject to attention, perception, and/or top-down cortical modulation as in the empirical recordings.

## Results

### Behavioral data

All speech tokens were perceived categorically by listeners (**Figure 2A**). Percent of /ga/ percepts for each token followed a stair-stepped identification function characteristic of categorical perception [*F*(4, 56) = 29.85, *p* < 0.001, *n*^2^_*p*_ = 0.68]. Consistent with the non-speech nature of the ambiguous stimulus, participants were unable to label the isolated base alone as either phoneme, resulting in chance level categorization. Critically, dichotic duplex stimuli were categorized with a similar label as their diotic counterparts; high F3 stimuli promoted the perception of “da” and low F3 stimuli the perception of “ga.” These results confirm listeners’ binaurally integrated speech cues support phonetic labeling. RTs were invariant across tokens [*F*(4, 56) = 1.41, *p* = 0.24, *n*^2^_*p*_ = 0.09] (**Figure 2B**) resulting in uniform decision speeds on the order of 400-450 ms.

**Figure 2:**
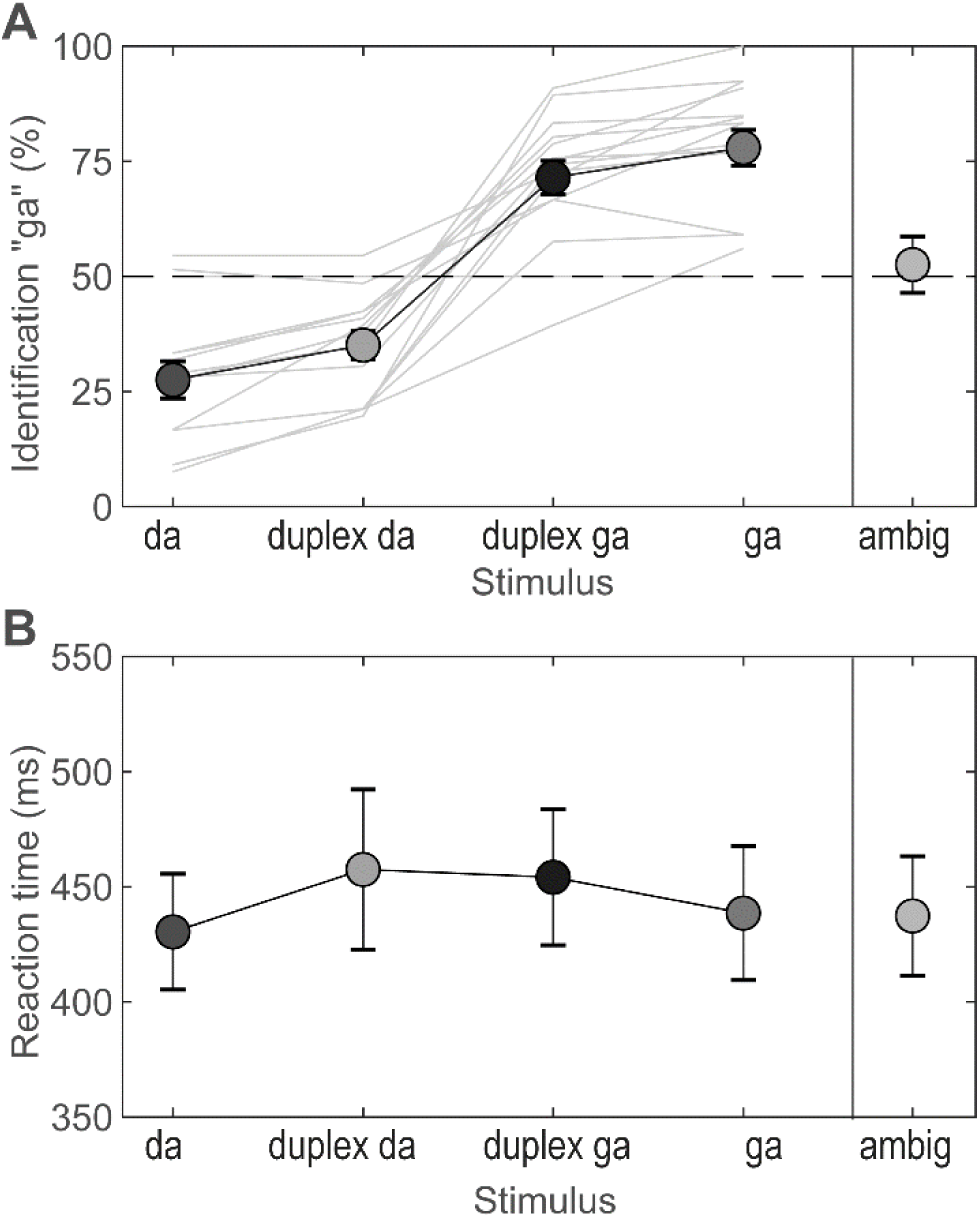
Behavioral identification of duplex stimuli shows categorical hearing of dichotic stimuli. (**A**) Identification follows a stair-stepped function characteristic of categorical perception. Bold line = grand average; light gray lines = individual listeners. Dichotic duplex stimuli (token 9 with high/low F3) were identified with similar labels as their diotic counterparts: “da” vs. “ga”, respectively. Ambiguous base stimuli, which do not carry category-relevant cues, were categorized at chance levels (dotted line). Da, duplex da, duplex ga, ga, and ambig tokens here correspond to the stimulus tokens 1, 9(highF3), 9(lowF3), 17, and ambig as described in Preisig & Sjerps (2019). (**B**) RTs did not vary across tokens. Error bars = ±1 s.e.m.

### FFR data

Grand average FFR time waveforms and response spectra for each token are shown in **Figure 3A-B**, respectively. Note the robust periodicity of FFR waveforms, reflecting phase-locked neural activity to both diotic and dichotic speech stimuli. Despite identical acoustics in the low-frequency portion of the acoustic stimulus spectrum (i.e., Fig. 1), FFR spectra showed prominent modulation in F0 strength across tokens, indicating perceptual influences on brainstem response magnitude (Carter and Bidelman 2023).

**Figure 3:**
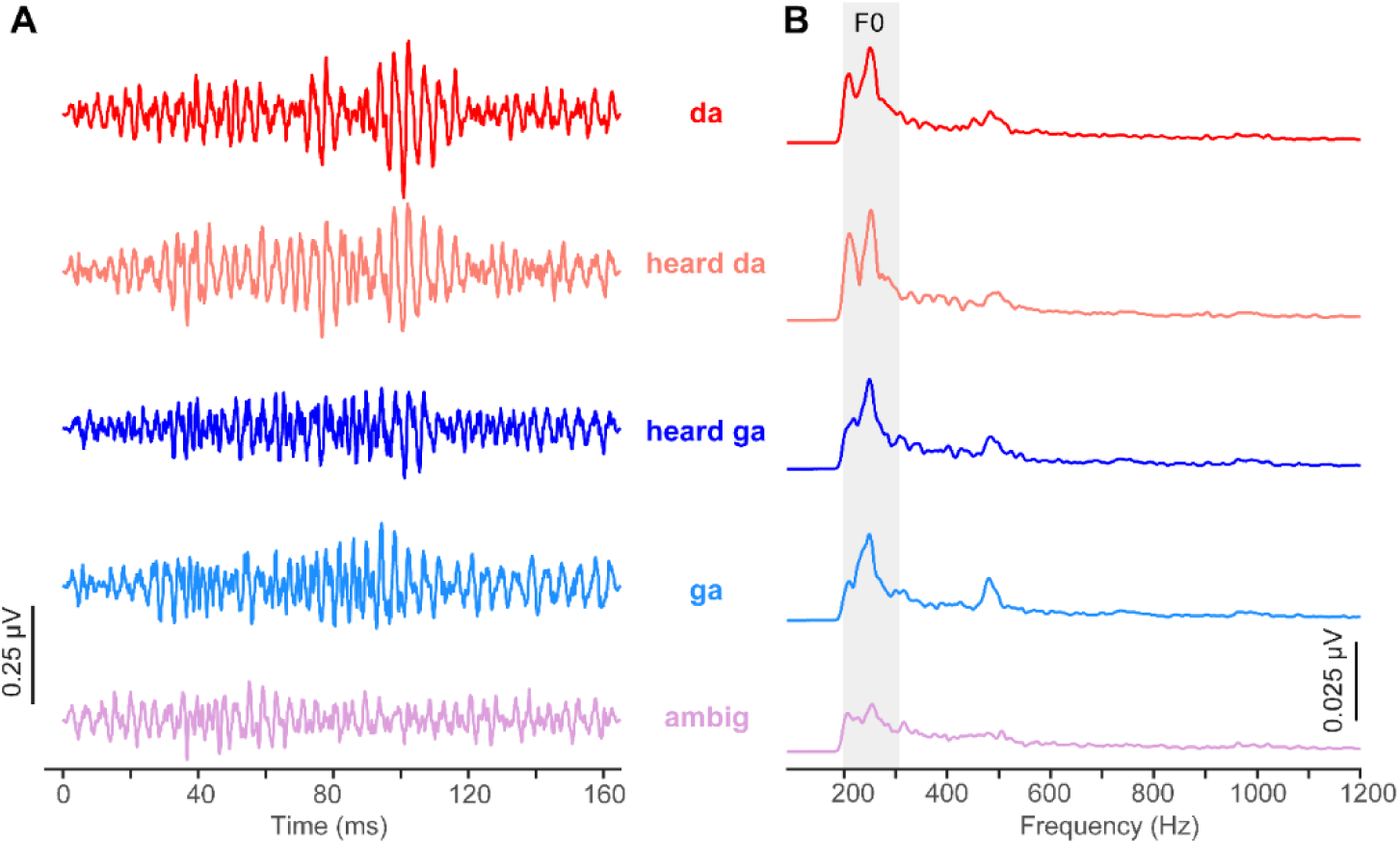
FFR waveforms (A) and spectra (B). Note the stronger response amplitude for sounds heard with a clear phonetic label vs. category-ambiguous speech sounds. Positive voltage is plotted up. Shaded area = region for F0 analysis.

Omnibus ANOVAs revealed a main effect of token on both FFR F0 amplitude [*F*(4,70) = 3.19, *p* = .018; *n*^2^_*p*_ = 0.15] and latency [*F*(4,56) = 3.17, *p* = .02; *n*^2^_*p*_ = 0.18] (**Figs. 4A-B**). Tukey-Kramer adjusted contrasts revealed the amplitude effect was driven by stronger responses to both the diotic (*p* = .062) and dichotic “da” (*p* = .004) stimuli relative to the ambiguous control. Similarly, the latency effect was driven by earlier responses to both /da/ stimuli compared to the ambiguous control (*ps* < .042). Responses were also faster on the whole to /da/ vs. /ga/ category stimuli (Student’s *t*-test: duplex/diotic /da/ vs. duplex/diotic /ga/ contrast; *t*(54.63) = – 2.24, *p* = .03). These findings suggest FFR latency distinguished stimuli from opposing perceptual categories regardless of whether they were constructed from monaural or binaural phonetic cues.

**Figure 4:**
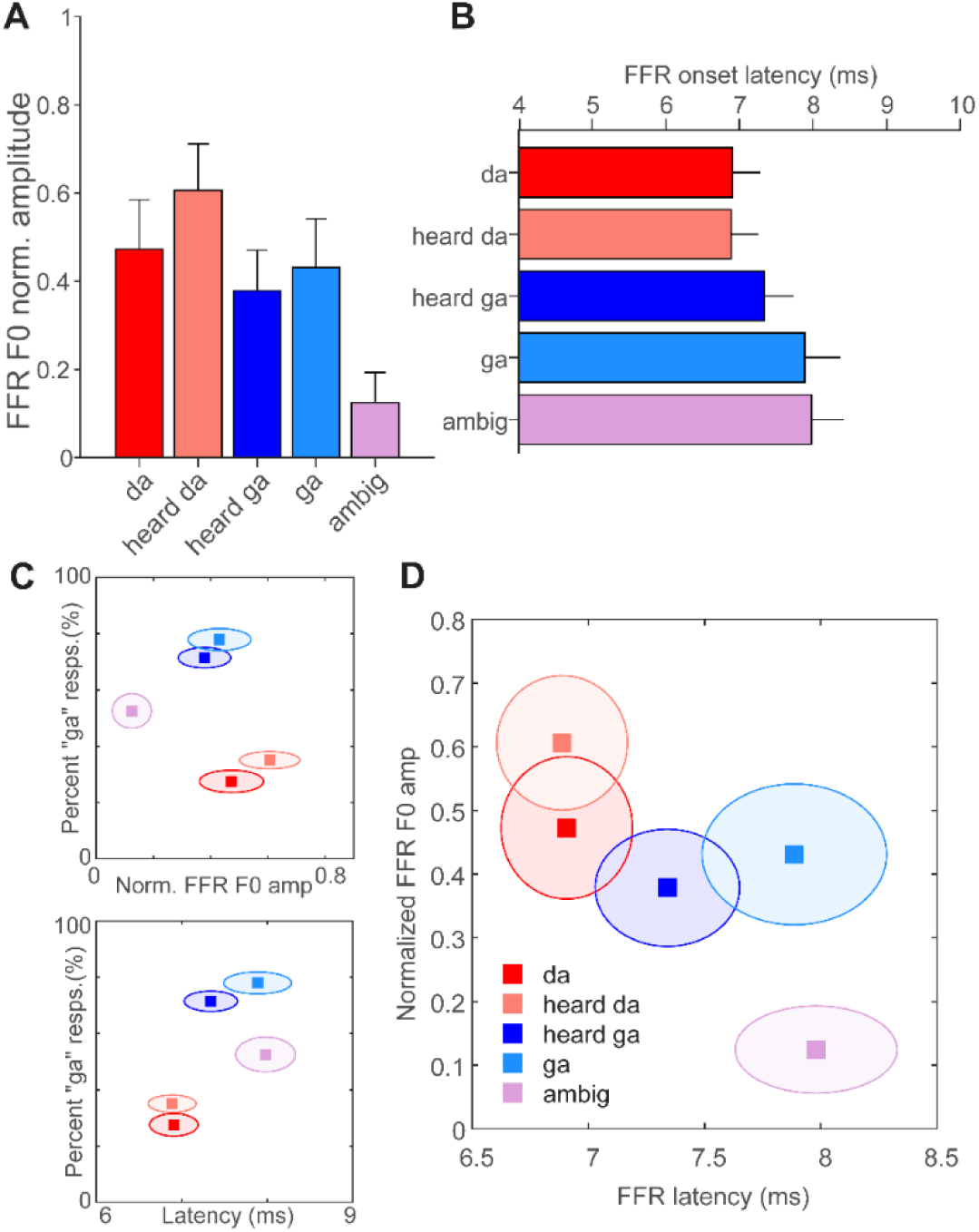
FFR F0 amplitude and latency to duplex stimuli reflect a differentiation of speech categories carried in either the acoustic (diotic) or perceptual (dichotic) domain. (**A-B**) FFR F0 amplitude and latency across token type. **(C)** Bivariate plots reveal perceptual categories separate from each other in perceptual-neural space. **(D)** FFR measures alone similarly show brainstem responses cluster according to their respective categories; neural responses to duplex syllables resemble their diotic counterpart. Error bars/shaded ellipses= 1 s.e.m.

To better visualize whether a combination of neural (FFR) and/or behavioral measures revealed category structure, we constructed a series of bivariate plots showing relationships between measures in perceptual-neural space (**Figure 4C**). We found perceptual speech categories “separate” from each other in perceptual-neural space (**Fig. 4C**). More critically, when considering FFR variables alone, the combination of neural amplitude and latency measures cleanly separated responses to /da/ vs. /ga/ stimulus classes in both dimensions (**Fig. 4D**). Stimuli perceived as “da” clustered with one another and vice versa for those perceived as “ga.” Moreover, responses categorically perceived as speech clearly segregated in multidimensional neural space from the ambiguous control stimulus. These findings indicate that FFRs not only differentiated speech compared to nonspeech sounds but more critically, clustered according to their phonetic identity even when their labeling required the fusion of binaural speech cues.

Brain-behavior repeated measures correlations revealed a positive relation between FFR latency and behavioral identification [*r_rm_*(44) = 0.38, *p*= 0.01] (**Fig. 5**). Later FFR latencies were associated with a greater preponderance of “ga” percepts within individuals. Correlations between F0 amplitude and PC were not significant.

**Figure 5:**
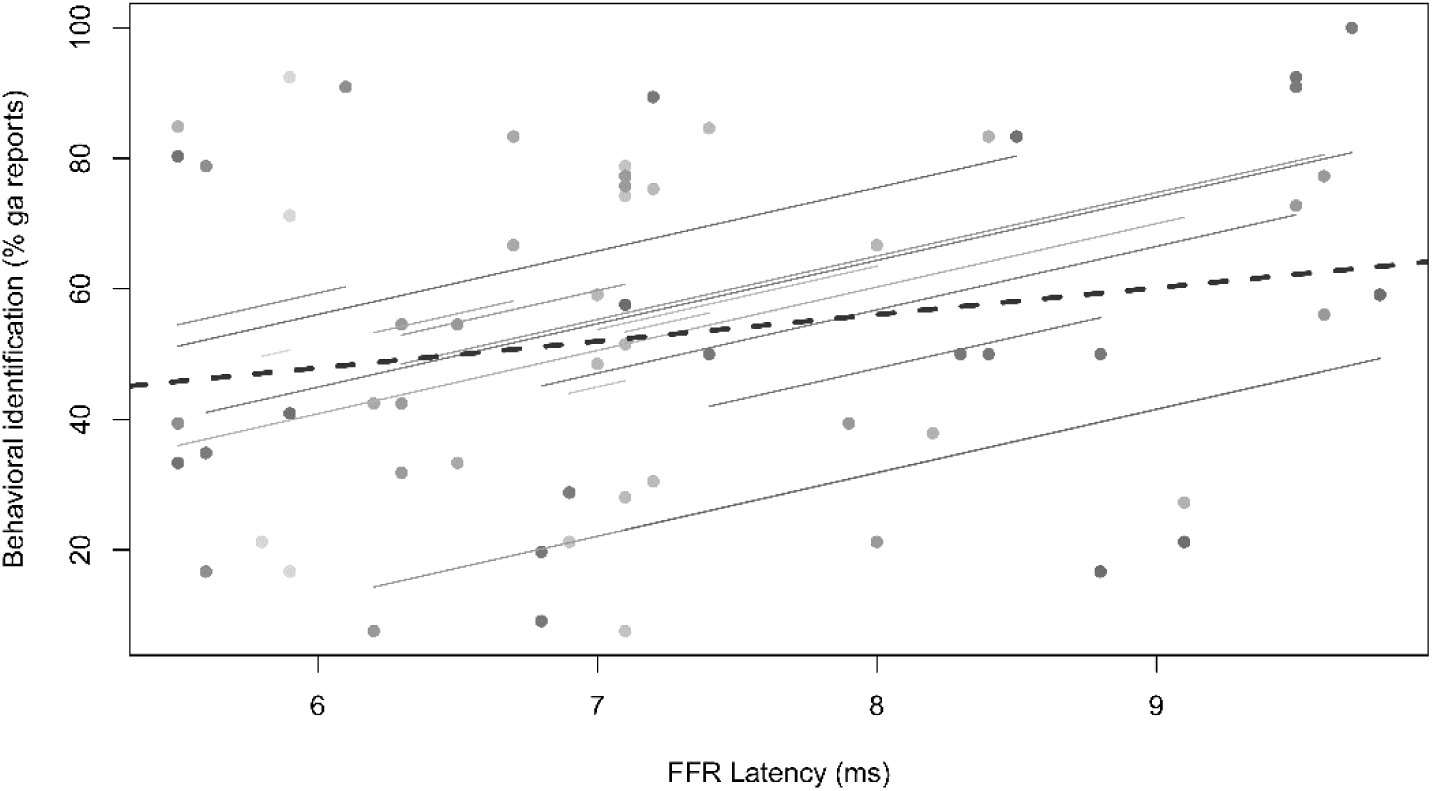
Brain-behavioral correlations reveal correspondence between FFR timing and categorical percepts. Repeated measures correlation (rmCorr) (Bakdash and Marusich 2017) reveals faster FFRs are associated with less frequent /ga/ percepts. Colored dots reflect each individual participants’ responses. Solid lines, within-subject fits to each individual’s data across the four stimulus conditions (ambiguous control not included); dashed line, fit across the aggregate sample.

**Figure 6** shows model dichotic FFRs to our duplex stimuli. In general, model F0 amplitudes were largely invariant across tokens. Critically, model FFRs did not show an enhancement for speech relative to non-speech (ambiguous) stimuli as in the empirical FFRs (cf. Fig. 4). Additional control analyses of the acoustic stimuli indicated that the F0 of our tokens was nearly identical, varying < 0.4±0.17 dB across conditions. Collectively, these findings support the notion that category coding effects observed in the FFR are not due to stimulus acoustics or cochlear nonlinearities, *per se*, but instead reflect top-down modulations from listeners’ perceptual-phonetic hearing of the speech stimuli (Carter and Bidelman 2023).

**Figure 6.**
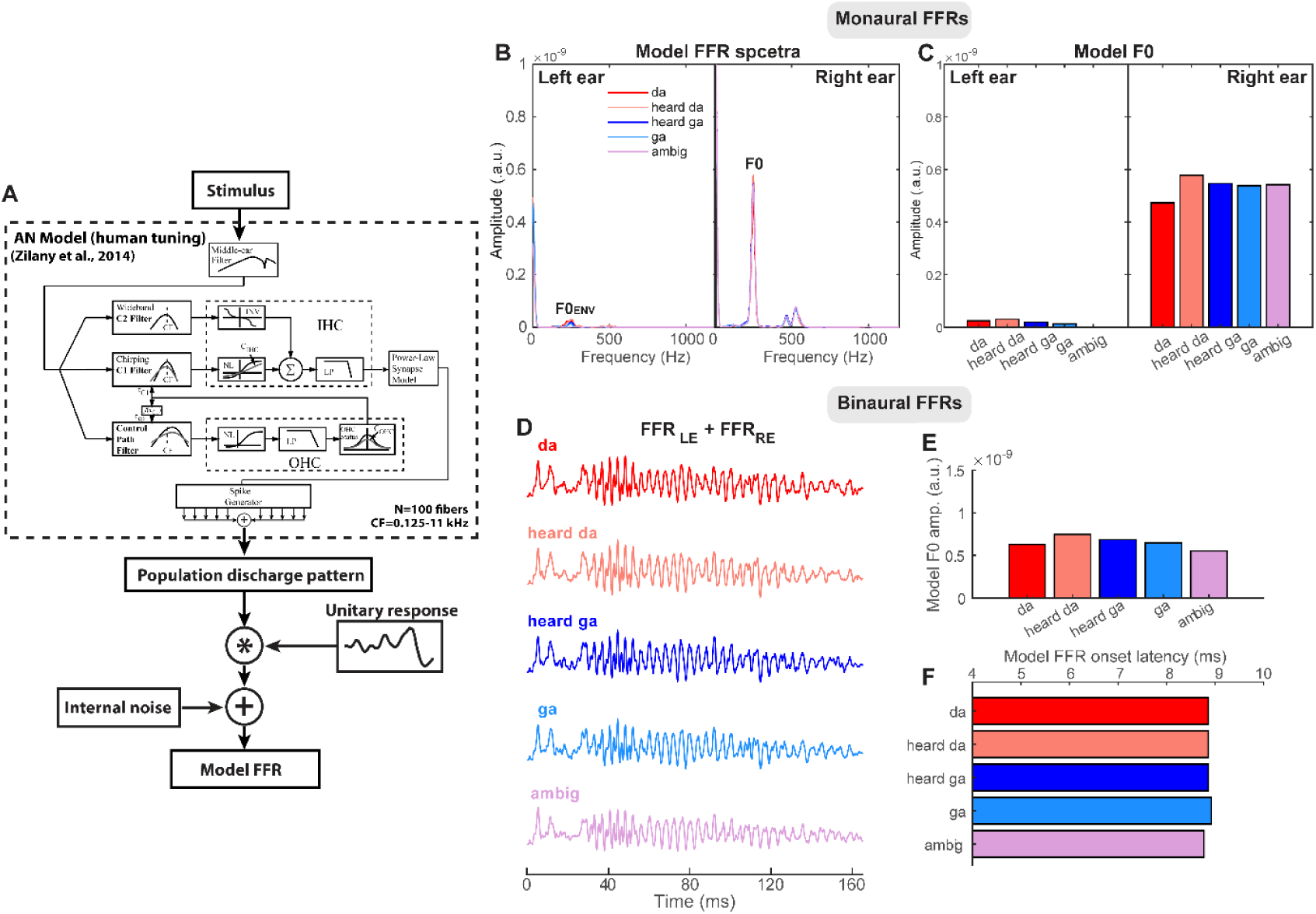
Computational model simulations of the scalp-recorded FFRs. (**A**) The acoustic stimulus is input to a biologically plausible model of the auditory periphery (Zilany *et al*. 2014) with human cochlear tuning (Shera *et al*. 2002). The model provides a simulated realization of the neural discharge pattern for single AN fibers. After middle-ear filtering and hair cell transduction and filtering, action potentials are generated according to a nonhomogeneous Poisson process. Spikes were generated from 100 model fibers (CFs: 125-11000 Hz) to simulate neural activity across the cochlear partition and summed to form a population PSTH for the entire AN array. Population PSTHs were then convolved with a unitary response function which simulates the impulse response of nuclei within the auditory brainstem (Dau 2003). Additive noise simulated the inherent random fluctuations in scalp-recorded EEG. (**B, C**) Model FFR spectra and F0 amplitudes for left (LE) and right ear (RE) channels of the duplex stimuli. Note the identical responses in RE owing to the same ambiguous base presented in all conditions. LE responses are considerably weak given that the high-frequency components of the left ear signal are well above the upper limit of phase-locking in the mammalian midbrain (Liu et al. 2006; Bidelman and Powers 2018). Though not present in the LE stimulus, the small F0 response observed in LE FFRs (F0_ENV_) likely reflects phase-locking in the low-frequency “tails” of high-frequency AN fibers to our high-intensity stimuli (e.g., Dau 2003). (**D**) Dichotic FFRs derived by summing model responses to each ear. (**E, F**) Model F0 amplitudes and latencies of dichotic FFRs are invariant and do not show the categorical variations as observed in the true FFR data (cf. Fig. 4).

## Discussion

We recorded FFRs evoked by duplex and diotic speech stimuli to explore how neural responses at the level of the auditory brainstem depend on listeners’ binaural integration and categorical perception. We found FFRs are stronger in response to speech (duplex and diotic tokens) compared to non-speech stimuli (ambiguous base), corroborating findings that brainstem representations are enhanced for behaviorally-relevant signals (Galbraith *et al*. 1995; Galbraith et al. 2004; Krishnan et al. 2005; Krishnan et al. 2009; Cheng 2021). Our data also reveal that FFRs represent binaurally integrated speech representations, reflecting a fused categorical percept. That binaural FFRs mirror listeners’ endogenously generated, behavioral report further bolsters the notion that FFRs carry more than a neuro-acoustic code and instead reflect listeners’ online perceptual state (Lai *et al*. 2022; Carter and Bidelman 2023; Lai *et al*. 2023). Our data further suggest category representations may emerge subcortically, prior to signal arrival in cortex. Categorical organization for speech at the brainstem level may therefore reflect top-down, corticofugal modulation of midbrain processing.

### FFRs are stronger in response to speech than non-speech sounds

We found that FFRs were enhanced to speech compared to non-speech (i.e., ambiguous) stimuli, corroborating previous studies. FFRs to forward speech are enhanced relative to FFRs to the same speech tokens time-reversed, indicating brainstem neural coding is enhanced when otherwise acoustically similar stimuli are perceptually relevant (Galbraith *et al*. 2004). Additionally, FFRs to sine-wave speech are enhanced for trained listeners who hear these stimuli as speech compared to naïve listeners who do not (Cheng 2021). Further evidence that FFRs are stronger for features in a listener’s native language supports the notion that subcortical auditory responses are enhanced for behaviorally relevant signals from a listener’s native language (Krishnan *et al*. 2005; Krishnan *et al*. 2009).

Our finding that speech FFRs were stronger to diotic and duplex speech tokens than to the ambiguous base alone corroborates findings that FFRs are enhanced for speech signals. Although we expected behavioral differences in RTs for duplex and ambiguous stimuli mirroring our FFR findings, the absence of this effect may have been due to the unique stimulus presentation paradigm with clustered presentation, allowing listeners to anticipate their responses, leading to faster, more uniform, reaction times across all tokens. That FFRs may be enhanced for relevant signals supports the theoretical notions that “speech is special” in terms of auditory processing (Liberman 1982; Liberman and Mattingly 1989). This position argues that speech signals are processed differently than acoustic information without linguistic value, which was an early topic of debate with duplex perception (Liberman *et al*. 1981; Liberman 1982; Mann and Liberman 1983; Whalen and Liberman 1987; Liberman and Mattingly 1989). Here, the enhancement of FFRs to speech tokens supports the idea that speech is afforded special processing in the brain by showing this privilege also extends to a subcortical level.

### FFRs reflect binaural integration

A novel finding here is that FFRs reflect binaural integration of duplex stimuli. Critically, our stimuli require binaural integration to be categorically perceived; listeners cannot arrive at a phonetic label without integrating speech cues from the two ears. The fact FFRs mirrored the behavioral reports supports the notion that FFRs carry information about binaural integration and reflect listeners’ online perceptual state (Lai *et al*. 2022; Carter and Bidelman 2023; Lai *et al*. 2023). Early binaural processing in the afferent auditory pathway begins in the superior olivary complex (Goldberg and Brown 1969), a site more caudal to the midbrain IC generators driving most of the FFR. As such, binaural integration should be measurable in auditory evoked potentials including the FFR. Indeed, binaural interaction has been observed previously in early waves of the scalp-recorded auditory brainstem response (ABRs) among components thought to arise from the lateral superior olive (Tolnai and Klump 2020). This implies that binaural integration is reflected at the brainstem level in neural activity well before signal arrival in the IC. IC neurons themselves are exquisitely sensitive to binaural inputs (Schreiner and Langner 1988). Since human FFRs reflect phase-locked activity predominantly from the midbrain and lower sites, the transformation of cues from both ears should be integrated prior to arriving at the IC. Indeed, several studies have demonstrated binaural interaction components in the FFR (Gerken *et al*. 1975; Hink *et al*. 1980; Krishnan and McDaniel 1998; Du et al. 2009). Additionally, animal studies have shown differential processing of temporal fine structure and envelope in FFR to changes in interaural time differences in rats (Xu et al. 2021), mirroring results reported in humans (Wang et al. 2018). Most literature examining effects of binaural integration on FFR focus on interaural cues (Ballachanda and Moushegian 2000). Here, the unique use of speech stimuli that require binaural integration for a categorical percept allows us a new perspective for examining binaural integration in FFR. An fMRI study using the same stimuli we used here revealed that BOLD activity in both auditory and nonauditory cortical areas predicted listeners’ perceptual reports (Preisig *et al*. 2022). Specifically, they showed differential activation dependent on listeners’ percepts in left perisylvian, inferior frontal and supplementary motor areas, and right motor and somatosensory cortices. The differences in activation here were independent of stimulus acoustics, indicating that categorical perception of these binaurally integrated stimuli activate a wide network of cortical regions. Our FFR findings further suggest that this network may also involve subcortical structures, earlier in the auditory system than previously thought^1^.

### Category representations emerge subcortically

We found that listeners’ categorical percepts of duplex stimuli modulated subcortical responses, suggesting category-level information is available to the brain prior to auditory cortex. Critically, the F0 (indeed all spectral cues) in our stimuli exceeded 250 Hz, which is substantially higher than the phase-locking limits of cortical neurons observed in any animal or human studies using either intracranial or far-field electrophysiological methods (Joris *et al*. 2004; Brugge *et al*. 2009; Bidelman 2018). Consequently, it is safe to conclude our FFRs and the categorical representations observed herein were of brainstem origin. Still, our data cannot adjudicate whether category-level representations are present in IC *de novo* (i.e., from bottom-up coding), or rather, emerge via top-down modulation of subcortical activity, e.g., via descending corticofugal projections that finetune brainstem auditory coding during active behavior (Suga 2008; Price and Bidelman 2021; Lai *et al*. 2022; Lai *et al*. 2023). Mirroring the percept-dependent modulations in our data, FFRs in response to sine wave speech in naïve vs. trained listeners provide additional evidence that corticofugal projections might modulate FFRs dependent on listeners’ percept (Cheng 2021). Further, our findings bolster those from Carter and Bidelman (2023), showing that speech-FFRs represent categorical information of the speech signal, presumably due to influence of top-down projections to midbrain. The corticofugal influences on FFR seen here are consistent with other studies suggesting that these modulations are strongest at F0, though for unknown reasons (Yellamsetty and Bidelman 2019; Price and Bidelman 2021; Lai *et al*. 2022; Carter and Bidelman 2023; Lai *et al*. 2023). The change in F0 amplitude seen here must be driven by more than merely stimulus acoustics, likely due to top-down changes of FFR, as the acoustic F0 remained the same frequency and intensity across tokens. Future studies are needed to test the constraints of top-down influences on FFR and to determine why categorical coding is most prominent at F0 when this component itself is not a category-defining cue.

Whether FFRs reflect an acoustic or perceptual correlate (i.e., exogenous vs. endogenous coding scheme) has been somewhat equivocal in the literature. Studies using speech-evoked FFRs to investigate perception have nearly always used passive listening paradigms (Aiken and Picton 2008; Skoe and Kraus 2010; Bidelman *et al*. 2013). This makes it difficult to establish a perceptual basis for the FFR as neural activity is not recorded during behavior and thus cannot be ascribed to representations beyond those that are merely sensory-acoustic in nature. However, more recent studies have employed novel listening paradigms where FFRs are recorded during active speech perception tasks (Price and Bidelman 2021; Carter and Bidelman 2023). Findings from those studies illustrate robust links between behavior and neural FFR responses, providing unequivocal evidence that FFRs reflect the perceptual state of the listener rather than a pure reflection of the stimulus acoustics (Price and Bidelman 2021; Lai *et al*. 2022; Carter and Bidelman 2023; Lai *et al*. 2023). Indeed, our data support the hypothesis that the FFR carries a more abstract, perceptual correlate of listeners’ intended behavior in addition to the lower-level, neuro-acoustic sound representation that is presumably more automatic in nature. Our use of duplex stimuli teases apart these dissociations between acoustic and perceptual FFR representations since stimulus acoustics and monoaural (peripheral) processing alone fail to account for our data (e.g., Fig. 6).

## Conclusions

We show brainstem FFRs were enhanced for behaviorally relevant speech signals and were modulated by listeners’ categorization of binaurally integrated speech cues. Our findings support notions that category representations are present in subcortical auditory processing. Such category organization in the FFR is presumably due to corticofugal efferent modulation of midbrain signal processing according to task demands and listeners’ trial-by-trial perception of otherwise identical acoustic signals (Lai *et al*. 2022; Carter and Bidelman 2023; Lai *et al*. 2023). We also explored binaural integration in the FFR and found enhancements in brainstem responses evoked by duplex stimuli (that carry a phonetic label) compared to those that are phonetically ambiguous. Collectively, the data show FFRs represent a surprisingly abstract level of the speech code often attributed to cortical levels and emphasize the dynamic, perceptually-relevant nature of subcortical auditory processing to complex perception.

## Funding

This work was supported by the National Institute on Deafness and Other Communication Disorders (R01DC016267 to G.M.B.).

## Acknowledgements

The authors thank Jessica MacLean and Jane Brown for comments on the early version of this manuscript. Requests for data and materials should be directed to G.M.B.

## Footnotes

1 Preisig, et al. (2022) actually did show activation in auditory midbrain regions during active but not passive phoneme identification tasks (see their Fig. 4), corroborating our electrophysical data. However, the potential that subcortical regions contribute to the categorical encoding of duplex speech stimuli was apparently not recognized or discussed in that paper.

## Notes

### Competing Interest Statement

The authors have declared no competing interest.

